# Iron homeostasis and reproduction: Unveiling the Microbiome-Gut-Brain-Axis connection in the mosquito *Anopheles culicifacies*

**DOI:** 10.1101/2025.05.31.657141

**Authors:** Pooja Yadav, Jyoti Rani, Tanvi Singh, Vaishali Saini, Pooja Rohilla, Vartika Srivastava, Gitanjali Tandon, Nirmala Sankhala, Gunjan Sharma, Suchi Tyagi, Sanjay Tevatiya, Seena Kumari, Rajnikant Dixit

## Abstract

Blood is an essential requirement for adult female mosquitoes to reproduce successfully. Communication between the Microbiome and Gut-Brain-Axis (mGBA) is crucial for maintaining organismal physiology during ovarian growth and development(*1*). Given the elevated iron levels in the blood, we investigated how mosquitoes maintain systemic iron homeostasis and reveals striking parallels to iron deficiency disorders in mammals. We demonstrate that the synergistic transcriptional regulation of *Ferritin (Fer)* and *Transferrin (Trf)* plays a crucial role in follicle development and egg maturation. Silencing both genes using RNA interference (RNAi) led to severe reproductive impairment, including ovarian arrest in 40% of females, a 50% reduction in oocyte numbers, and reduced first instar larval size. These e phenotypes correlate with increased reactive oxygen species (ROS) and altered serotonin (5-HT) receptor expression in the brain, likely driven by gut microbiota imbalance, and hence neurological signalling modulation due to disrupted iron metabolism. The unexpected discovery of a transcript encoding a novel hepcidin (*AcHep*) protein that was previously unknown in any insect or mosquito species, and surprisingly, its modest enrichment in the fat-body of the *Fer+Trf* knockdown mosquito, suggested that it may be involved in the maintenance of the systemic iron homeostasis, as in mammals. Summarily, our data provide the first molecular evidence in insects that iron metabolism disorders can impair mGBA communication and compromise reproductive outcomes, mirroring effects observed in mammalian iron deficiency. This study expands our understanding of iron’s systemic roles and opens new avenues for targeting mosquito reproduction in vector control strategies.

## Introduction

Maintaining a basal level of body iron is crucial to regulating many physiological processes, such as DNA synthesis, repair, immunity, and oxidative activities(*2–4*). Cycling of two iron forms, i.e., reduced ferrous (Fe^2+^) or oxidized ferric (Fe^3+^) between two distinct oxidation states, is an important aspect of iron homeostasis in any biological system(*5*). Ferrous iron is highly soluble in many physiological environments and forms harmful toxic radicals, in contrast to ferric iron. A misbalance of iron concentration may have a deleterious impact on the health of an organism, and therefore the diverse nature of iron-binding proteins participates in managing optimal iron homeostasis(*6*, *7*).

Hematophagous insects consume large amounts of iron-rich blood meal for their ovarian follicle growth and egg development(*1*). After blood-feeding rapid establishment of a gut-brain-axis communication *via* gut-microbiome is crucial to maintaining organismal physiological homeostasis(*1*), but the molecular nature and function of key factors controlling mobilization, storage, and utilization of blood-derived iron in mosquitoes are still obscure. Because adaptation to the blood-feeding is a high-risk, high-reward process, and iron serves many physiological processes, a multifactor protective mechanism has been suggested to avoid oxidative stress in blood-feeder insects(*8*).

Blood contains ∼10 mmolL^−1^ of heme-bound iron(*9*) and like humans, mosquitoes also encode transferrin and ferritin proteins as an integral part of several metalloproteins, likely to deal with the systemic heme bound iron concentration and reduce oxidative stress(*4*, *10*). Though export mechanisms for heme and ferritin-bound iron are undefined yet, a mechanistic overview suggests the efflux of ferrous ions follows oxidation to ferric ions, which are then bound to serum transferrin for transportation to other cells(*8*).The absence of the mammalian homolog *ferroportin,* which exports ferrous ions as a substrate for oxidation *via* ferroxidase, is still an unaddressed problem in mosquitoes(*11*). In the tick *Ixodes ricinus*, silencing of intracellular ferritin (*fer-2*), which also carries a conserved ferroxidase site, and iron-regulatory protein (*irp1*), help in transferring non heme iron, significantly impairs the reproductive outcome(*12*).

Recently, we demonstrated that fat-body expressing Transferrin (*Trf*) plays a major role in iron transportation, and mRNA depletion of this transcript significantly impairs oocyte development in the mosquito *An. culicifacies*(*13*). While another vertebrate’s homolog, metalloprotein ferritin, has also been identified as a secretory protein from several insects(*14*, *15*)found to act as an iron storage protein. Though most studies highlight its role in insect immunity(*16–18*) few studies also suggest that increased ferritin-stored iron in the ovary after a blood meal may be required for an insect’s embryonic and larval development(*19*, *20*). However, its functional correlation with transferrin and its role in regulating iron homeostasis and reproductive physiology in mosquitoes remain unexplored.

Interestingly, emerging studies indicate that gut-associated microbes not only aid in the digestion of iron-rich blood meals but also play key roles in managing iron-induced oxidative stress, immunity, and vectorial competency(*21–25*). A heightened expression of gut-bacterial encoded transcripts, *viz*. TonB, FecR after blood-feeding correlates with their possible role in iron metabolism in the mosquito *An. stephensi*(*26*), but the mechanism of safe utilization of excess free iron is unclear. Studies in humans highlight several bacterial species, such as *Aeromonadeceae*, that need iron for multiplication to cause pathogenicity, which is achieved either by destroying erythrocytes and hydrolyzing the hemoglobin, or sequestering iron from ferritin(*3*, *27*).

Because blood meal boosts the rapid proliferation of gut bacteria, we hypothesize that a gradual shift between mosquito-derived iron mobilizing factors e.g., ferritin, transferrin, and the iron-utilizing gut-bacterial population, could be a key mechanism to minimize and nullify the harmful effect of unbound toxic iron-free radicals. Conceptual learning supports the idea that free iron originating from the gut during blood meal digestion is sequestered in the haemolymph, and *Fer+Trf* may help in the storage and transport of this iron to the ovaries for egg development in the gravid female mosquitoes(*8*). In contrast, a substantial amount of unutilized iron may still be available for microbial growth in the gut. Thus, it is plausible to propose that a functional disruption of systemic iron homeostasis may have negative consequences on the microbiome-mediated gut-brain-axis communication, and hence the reproductive physiology of the mosquitoes.

To test and evaluate the hypothesis, here we identify, characterize, and establish a functional correlation of Ferritin and Transferrin in iron storage and transportation, and explore their possible role in the reproductive physiology of the mosquito *An*. *culicifacies.* We demonstrate that a synergistic transcriptional modulation of *Fer* and *Trf* is essential to mobilize the iron in the ovarian maturation of the gravid female mosquitoes. We highlight that disrupting systemic iron homeostasis by KD of *Fer* and *Trf* alters the gut-microbial community structure, and hence increases ROS activities, which not only harms brain functioning but also severely impairs ovarian follicle and egg development. With the novel discovery of hepcidin-like peptide, a well-known negative regulator of mammalian iron homeostasis, we propose that, like mammals, the mechanism of mobilization and utilization of heme-derived systemic iron content for ovarian maturation and egg development functionally remains conserved in mosquitoes.

## Methodology

### (1) Mosquito rearing and maintenance

*An. culicifacies* (sibling species A) was reared and maintained in the ICMR-National Institute of Malaria Research’s central insectary facility under conventional rearing conditions of 28 ±2°C temperature, 60-80% relative humidity, and a 12:12 hr light/dark cycle(*1*, *3*, *28*). All protocols for the cyclic colony of mosquito rearing and maintenance of the mosquito culture were approved by the ethical committee of the ICMR-National Institute of Malaria Research, New Delhi (NIMR/IAEC/2017-1/07).

### (2) Exogenous ferric chloride feeding assay

A control group (100 mosquitoes) of newly emerged mosquitoes was given only a 10% sugar solution, whereas test groups (100 mosquito) received sugar with ferric chloride at varying concentrations (1 mM, 5 mM, and 10 mM). The sugar solution and the supplemented meal were given through cotton swabs and changed after 24 hours with a fresh meal. Survival of the mosquitoes was monitored every 24 hours of iron supplementation. Haemolymph, fat body, midgut, and ovary were collected from surviving 20–25 mosquitoes from each set (control and supplemented test group) after 2 days of supplementation to assess the transcriptional response of target transcripts.

### (3) Mosquito tissue sample collection

As per the experimental design, the desired tissues were collected from 4-day-old adult female *An. culicifacies* mosquitoes. For this purpose, ice-anesthetized mosquitoes (20 mosquitoes) were placed on slide under the dissecting microscope one by one, and dissected with the help of a sharp needle. Different tissues including the salivary gland, midgut, fatbody, hemolymph, and female reproductive organ were dissected and collected in Trizol reagent as described previously(*3*). The aquatic development stages, such as egg, larvae, and pupa of the *An. culicifacies* were also collected in Trizol reagent. To explore the fate of iron, we focused on relevant tissues such as the haemolymph, fat-body, mid-gut, and ovary collected from both naive sugar-fed and blood-fed mosquitoes of the same cohort at pre-defined time interval. Initially, targeted tissues were collected from sugar-fed mosquitoes aged 3-4 days, while to meet the blood-fed condition, a live and healthy rabbit was placed inside a mosquito cage for blood-feeding. Then selected tissues were collected from a fully engorged female mosquitoes at 0-2 hours, 24 hours, 48 hours, and 72 hours after the blood meal for detailed tissue specific gene expression analysis. For hemolymph collection, an anticoagulant solution containing (60% Schneider’s medium, 10% foetal bovine serum, and 30% citrate buffer) was injected into the thorax, and a small incision was made near the posterior two segments of distended belly with a minuscule needle, allowing translucent hemolymph to ooze out. Using a micropipette, hemolymph was taken and pooled in Trizol. Fat-body was recovered by extracting all abdominal tissues from the last two abdominal segments and then forcibly tapping the abdominal carcass till pale yellow fat-body oozed out.

### (4) Total RNA isolation and cDNA synthesis

Total RNA was isolated from collected samples manually by Trizol method, as previously described(*28*). Isolated RNA was quantified with the help of a Nanodrop 2000 spectrophotometer (Thermo Scientific, USA), and ∼1 μg of total RNA was used for the synthesis of the first-strand cDNA using the cDNA synthesis kit (PrimeScript^TM^ 1^st^ strand cDNA Synthesis Kit, Cat. # 6110A, Takara). Actin was used as a reference gene in RT-PCR to evaluate the quality of the cDNA.

### (5) Gene expression analysis through Quantitative PCR

For gene expression analysis, a real-time PCR-based transcriptional profiling assay was performed using the Sybr Green qPCR Mastermix (TB Green Premix Ex Taq^TM^ II, Cat# RR820A, Takara) and CFX 96 Real-Time PCR system (C1000 Touch Thermal Cycler, Bio-Rad, USA). Three independent biological replicates were run under the identical PCR conditions, starting with denaturation at 95°C for 5 min, 40 cycles of 10 sec at 95°C, 15 sec at 52°C, and 22 sec at 72°C. Each cycle’s fluorescence reading was recorded at 72°C after it had finished. For determining a melting curve, the final PCR stages at 95°C for 15 sec, 55°C for 15 sec, and again 95°C for 15 sec were accomplished. Throughout the study, the Actin and Rps7 genes served as reliable internal control (reference gene), which were found with stable expression throughout different stages and tissues of the mosquito(*29*). The relative quantification was examined using the 2^−ΔΔ^Ct method.

### (6) dsRNA-mediated gene knockdown Assays in adult mosquitoes

A double-stranded RNA-mediated gene knockdown assay was executed to suppress target gene mRNA expression. Single-stranded cDNA was first amplified using a dsRNA primer with a T-7 overhang primer as listed in the Supplemental Table 1. Following amplification, the PCR product was purified through a Gene jet purification kit (Gene JET PCR Purification Kit, Cat #K0701; Thermo Scientific, USA) and quantified by Nano-drop 2000 spectrophotometer (Thermo Scientific, USA). Using the transcript aid T-7 high yield transcription kit, (Cat# K044, Ambion, USA) dsRNA was *in-vitro* synthesised by employing the purified PCR product as a template. The resulting reaction product was again purified, quantified, and approximately 69 nl (∼3 ug/ul) was injected into the thoracic region of a 1-2 day old cold-anesthetized mosquito using a nano-injector (Drummond Scientific, USA, Cat# 13681455). Concurrently, for the negative control group dsRNA of bacterial-GFP was injected into the mosquitoes of the same cohort. After 48hr of post injection, tissue samples were dissected from both the gene-specific i.e. Ferritin only and in combination of Transferrin (Trf), as well as control (GFP-injected) injected mosquitoes. To determine the silencing effectiveness, a real-time PCR-based assay was performed as described above.

### (7) Assessment of ovary development following gene silencing

Both the control (GFP dsRNA-injected) and experimental (*Fer + Trf* dsRNA-injected) mosquito groups were fed on the rabbit, after 48hr of post injection and only fully fed mosquitoes were selected for the evaluation of ovary development, while partially fed or unfed were excluded. To evaluate ovary growth, anaesthetized mosquitoes were placed on dissecting slides under a binocular stereomicroscope and detached the ovaries in phosphate-buffered saline (PBS) at 72 hours post blood feeding. The mature oocytes inside ovary were manually counted, and the results were compared to control mosquitoes. However, gradual follicular development was compared within naïve sugar-fed, 24 hr, 48 hr, and 72 hr blood fed mosquito ovaries. Dissected ovaries from anesthetized mosquitoes were treated with PBS-T (Phosphate Buffered Saline with 0.5% Triton X-100) solution for 2-3 minutes followed by thorough washing with 1X PBS. The samples were then incubated for 5 min. with DAPI (40,6-diamidino-2-phenylindole, 300 nM). After staining, extra stain was removed by washing three times with 1X PBS in the dark, and tissue was imaged at excitation/emission: 358 nm/461 nm under a Leica confocal microscope (Thermo-fisher, USA)(*30*).

### (8) Metagenomic profiling of mosquito gut microbiota by the Whole Genome Shotgun method

For WGS-metagenomic analysis, we collected gut from 24 hours of blood-fed control and *Fer+Trf* KD adult female mosquitoes. Before dissection, the body surface of the mosquitoes was sterilized using 70% ethanol for 1 min. (5 mosquitoes per batch), followed by aeration on a sterilized filter paper. The gut dissection was carried out in 1X PBS, under the laminar flow, where the working area as well as the dissecting stereomicroscope were kept sterilized by using 70% ethanol. We followed a sample pooling strategy, which provides a reliable and comprehensive overview of the gut-bacterial community structure, if aimed at comparing within similar samples(*31*). For the purpose, pooled guts (50 whole midgut) were collected into a minimal volume (100ul) of sterile ice-cold 1X PBS and outsourced for WGS sequencing. High-quality genomic DNA was extracted from the collected sample. After isolation it was quantified and evaluated for quality and integrity. Library construction included appropriate adaptor ligation with DNA fragments, followed by amplification and sequence alignment (https://biologyinsights.com). The raw fastq reads were pre-processed using Fastp v.0.20.1(parameters: --length_required 50 -- qualified_quality_phred 30 -- unqualified_percent_limit 30 --average_qual 30)(*32*). Read-level taxonomic profiling was done by aligning the processed paired-end reads to the pre-KMA-indexed NCBI 2019 Genome Build (http://dx.doi.org/10.25910/5cc7cd40fca8e) database using KMA(*33*). The resulting KMA files were subjected to metagenomic profiling using CCmetagen v.1.2.5 and visualized using krona (CCMetagen-1.2/)(*34*) and stacked bar plots were plotted using phyloseq R packages(*35*).

### (9) Oxidative stress measurement assays

To assess the impact of iron homeostasis disruption on oxidative stress-mediated alteration in the midgut and brain, we performed molecular, biochemical, as well as cellular assays as described below.

#### Molecular assay

The brain was dissected from 15 mosquito of both KD and control group in order to assess the expression of genes that respond to oxidative stress (NOS and SOD). Brain samples were collected 48 hours after silencing, for total RNA isolation, cDNA preparation, and gene expression evaluation of NOS in the naïve sugar-fed mosquito’s brain *via* real-time, as described above. Likewise, following 72 hour of dsRNA injection, both control and KD mosquitoes were offered blood meal, and after 24 hours of blood-meal, the expression level of SOD were examined in the gut.

#### Cellular Assay

Oxidative stress in naïve sugar-fed (SF), blood-fed (BF) control, and *Fer+ Trf* KD mosquito groups was evaluated through ROS assay. Dissected brain tissue from 15 mosquitoes of each group were incubated with an oxidant-sensitive fluorophore dye CM-H2DCFDA [5-(and-6)-chloromethyl-29,79-dichloro-dichlorofluorescein diacetate, acetyl ester, 2mM] (Sigma, St. Louis, MO, USA), for 20 min at room temperature under dark conditions. To remove the extra staining, tissue was washed thrice with 1X PBS and observed under a Leica confocal microscope, observing at excitation/emission: 480nm/530nm to detect ROS accumulation.

#### Biochemical Assay

Alternatively, to estimate the oxidative stress, we collected 15 mosquitoes’ brain and homogenized them in 1X PBS for a very short period, and then briefly centrifuged to extract the supernatant for nitrite quantification using the Griess Reagent Kit (Cat. # G792, Thermo Fisher Scientific, USA). N-(1-naphthyl) ethylenediamine dihydrochloride (Component A) and Sulfanilic acid (Component B), were mixed, followed by the addition of 130ul of the mixture, 20ul of deionized water, and 150ul of the test sample as per kit manual. The final mixture was loaded in a 96-well U-shaped ELISA plate and incubated in the dark for 30 minutes. While 20 µL of Griess Reagent and 280 µL of deionized water mixture were considered as a reference sample, which was also loaded onto a plate, and absorbance was recorded at 548 nm using an ELISA Reader (Tecan Unlimited M200pro Instrument)(*36*).

### (10) Statistical analysis

For statistical analysis, test sample data were compared with the control data set using GraphPad Prism 9.5 software, and treatment differences were determined through one-way ANOVA test. All experiments were conducted thrice for data validation.

## Results

### Ferritin & transferrin exhibit synergistic transcriptional regulation during ovary maturation and egg development

A comprehensive literature review highlighted that insect Ferritin comprises two subunits i.e. heavy chain (HC) and light chain (LC)(*15*). Finding two putative transcripts encoding insect homologs HC and LC of the Ferritin protein from the hemocyte RNAseq data(*37*) prompted us to evaluate their role in iron metabolism in the mosquito *An. culicifacies*. Initial transcriptional profiling data suggested *Ferritin heavy chain (Fer)* might have a key role in iron storage properties, supporting previous findings, and hence we targeted only *Fer (HC)* in our investigation (Supplementary Fig. 1). Structure prediction and modelling analysis showed that *Trf* and *Fer* proteins carry distinct functional domains to bind iron; We also noticed an overlapping behaviour in iron-binding of ferrous to ferric status, which may depend on the availability of free iron at a particular physiological status. It was observed that ferrous ions had a greater binding affinity with ferritin, while ferric ions had a greater affinity towards transferrin. Interestingly, while focusing on specific binding sites or functional domains for both *Fer* and *Trf* behaved differently for ferric and ferrous ions (Supplementary Fig. 2).

A constitutive expression of *Fer* transcript was readily observable throughout all developmental stages, notably, we spotted an enriched level in eggs and pupae, a pattern previously noticed for *Trf*(*3*). These findings suggested that eggs and pupae are metabolically hyperactive than other developmental stages for iron mobilization and utilization. We also observed that the relative expression of *Fer* was higher in adult females compared to age-matched male mosquitoes (Fig.1A), signifying adaptive evolutionary benefits to females for an iron-rich blood meal and gonotrophic success. Corroborating with the dominant expression of *Trf* and *Fer* in the fat body and hemolymph, respectively (Fig.1B), and an elevated expression of *Fer* in the naïve adult female reproductive organ (Supplementary Fig. 3A), together suggested that a synergistic relationship is crucial to oocyte development in the female mosquito.

**Fig. 1:**
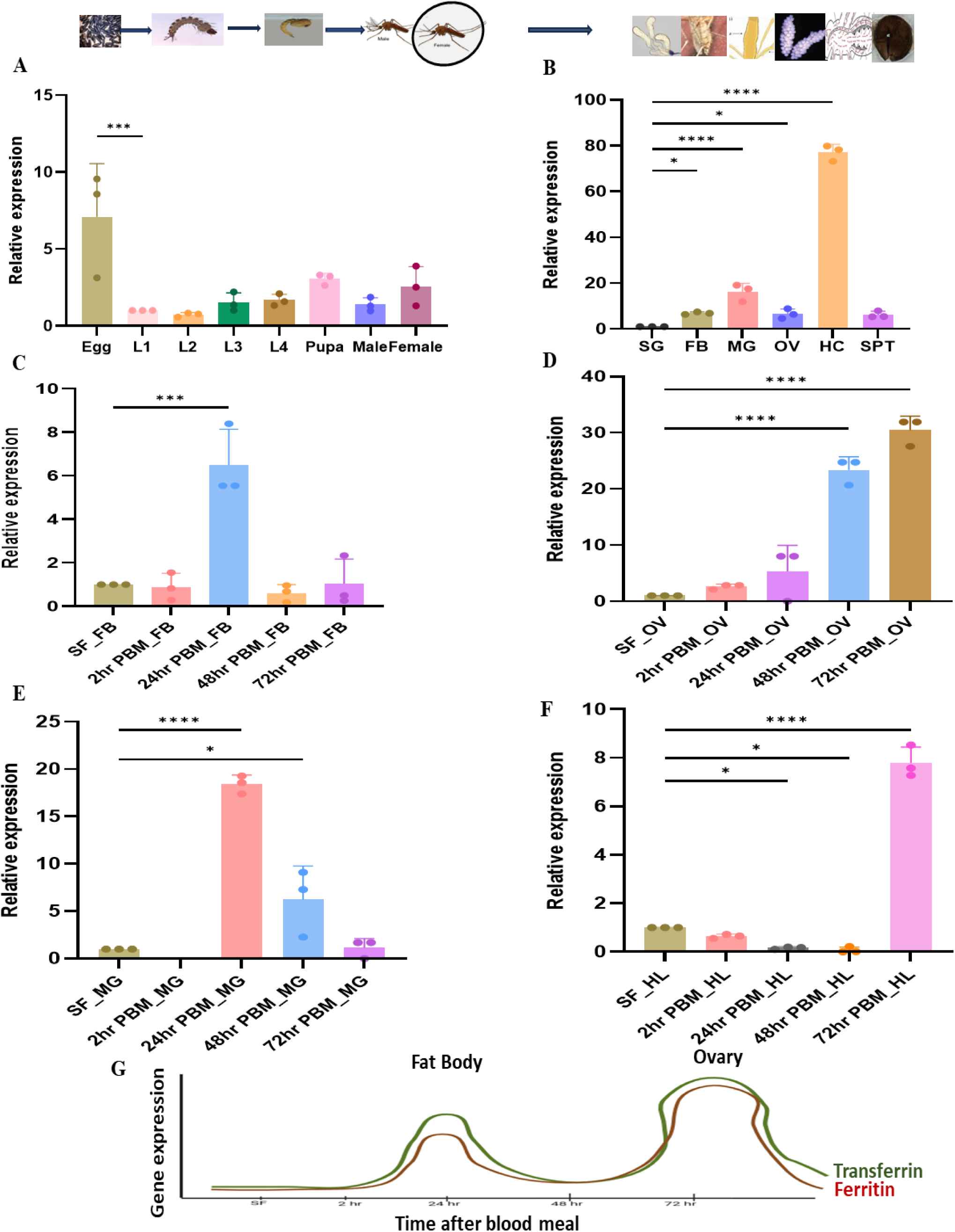
Spatio-temporal modulation of *Fer* expression in the *An. culicifacies* mosquito. **(A)** Relative *Fer* expression during development, showing increased expression of *Fer* in the egg (*p*<0.0003) of the mosquito. Here, the first instar larva was considered as a control for all test samples. (N=3, n=10); **(B)** Tissue-specific expression analysis showing high abundance in hemolymph (*p* <0.0001), midgut (*p* <0.0001), fat-body (*p* <0.0373), ovary (*p* <0.0483), and spermatheca in decreasing order, considering the salivary gland as a control for all test samples. (N=3, n=25); **(C)** FB, up-regulated expression at 24 hours of blood meal (*p* <0.0001). **(D)** OV, highest expression at 72hr of PBM (*p* <0.0001) Tissue-specific post blood meal time-dependent expression in **(E)** MG, increased expression at 24hr of PBM (*p* <0.0001) **(F)** HL, abundant expression at 72hr of PBM (*p* <0.0001) considering SF taken as control for all test samples. (N=3, n=25). All three independent biological replicates were statistically analyzed with the help of one-way ANOVA (Dunnett’s multiple comparison test), where asterisks represented *p < 0.05; **p < 0.005; and *** p < 0.0005. (N = number of biological replicates, n= number of mosquitoes dissected for sample collection). **(G)** Graphical overview of synergistic expression for Ferritin and Transferrin in Fat-body and Ovary. SG-Salivary gland, MG-Midgut, FB- Fat-body, OV- Ovary, HL- Hemolymph, SPT- Spermatheca, SF- Sugar fed.

To strengthen the proposition of *Fer* and *Trf* synergism, we evaluated the effect of blood meal on the *Fer* expression pattern. A time-dependent transcriptional response of *Fer* was monitored in various tissues, including hemocytes, midgut, fat body, and ovary, after blood feeding. Within 24 hours of blood-meal, a significant enrichment was observed in the midgut (Fig.1E) and fat body (Fig.1C), which gradually declined to baseline after 72 hours. A progressive increase of *Fer* transcript was observable in the ovary up to 72hr (Fig.1D), whereas, in hemolymph it was induced only after 72hr following blood meal (Fig.1G). We noticed this expression pattern of *Fer(F)* well aligns with *Trf(T)* expression in the ovary, hemolymph, and fat body after a blood meal(*3*). Simultaneously, we also witnessed the higher expression of *Heme oxygenase (HO)* transcript at 24 hours post blood meal, pointing fast release of substantial ferrous ions in the gut(*38*, *39*) which may likely be converted to ferric ions by the *Fer* in the midgut (Supplementary Fig. 4A). We hypothesized that a synergistic and biphasic regulation of *Fer* and *Trf* among midgut, hemolymph, fat body, and ovary could be crucial to meeting the iron requirement in oocyte maturation and egg development in gravid female mosquitoes.

### Effects of iron toxicity on mosquito survival

Leftover unutilized iron may cause toxicity within the mosquito body by creating free radicals, which could be detrimental to the mosquito’s health(*40*). An exogenous iron supplementation test with varying concentrations (1mM, 5mM, and 10mM) was carried out to evaluate the mosquito’s survival. Freshly emerged mosquitoes were offered different iron concentrations through sugar solution(*3*). Compared to the control, we noticed that after 24 hours of supplementation ̴ 90% of the mosquitoes kept on the 10mM iron solution died, whereas the survival rate of mosquitoes supplemented with 1mM iron solution was similar to the control group mosquitoes kept on standard sugar (Supplementary Fig. 3D). Surprisingly, we observed a gradual enrichment of *Fer* expression in the hemolymph with increased iron concentration (Supplementary Fig. 3B), while in the ovary its expression was moderately increased at 1 mM iron concentration and remained static even at higher concentrations (Supplementary Fig. 3C). Previously, we have observed that 1mM iron supplementation not only elevates the expression of transferrin in the fat body but also enhances the oocyte number in the gravid females(*3*). These findings imply that increased exogenous iron concentration may induce ferritin as well as transferrin levels, possibly to bind free iron and reduce its toxicity for survival.

### Dysregulation of Ferritin and Transferrin impairs reproductive potential

Recently, we demonstrated that *Trf* significantly contributes to shaping the mosquito reproductive cycle(*3*), but whether ferritin also plays a role in reproductive physiology remains untested(*41–43*). To test it, we carried out a gene knockdown experiment and evaluated its effect on the egg-laying capacity. Relative to the control (*dsGFP* injected) mosquito groups, we observed at least 90%, 60%, and 50% reduction of *Fer* transcript in the hemolymph, fat body, and midgut of the *dsFer* injected mosquito group (Fig. 2A), respectively. To evaluate the possible impact on reproductive outcome, we offered a blood meal to both control and KD mosquito groups. Following a 72 hours of blood feeding, we dissected the ovaries and manually counted the number of oocytes. Compared to the average number of oocytes in the control (69 oocytes), a significant reduction (44 oocytes) in the ovaries of the *Fer* KD mosquito (Fig. 2C), highlighted that *Fer* also influences the reproductive outcome. Surprisingly, we also observed that decreased *Fer* expression significantly alters transferrin expression by upregulating in the hemocytes (Fig. 2B), confirming that the participation of both proteins is essential in iron mobilization, possibly to regulate optimal systemic iron homeostasis maintenance during ovary development.

**Fig. 2:**
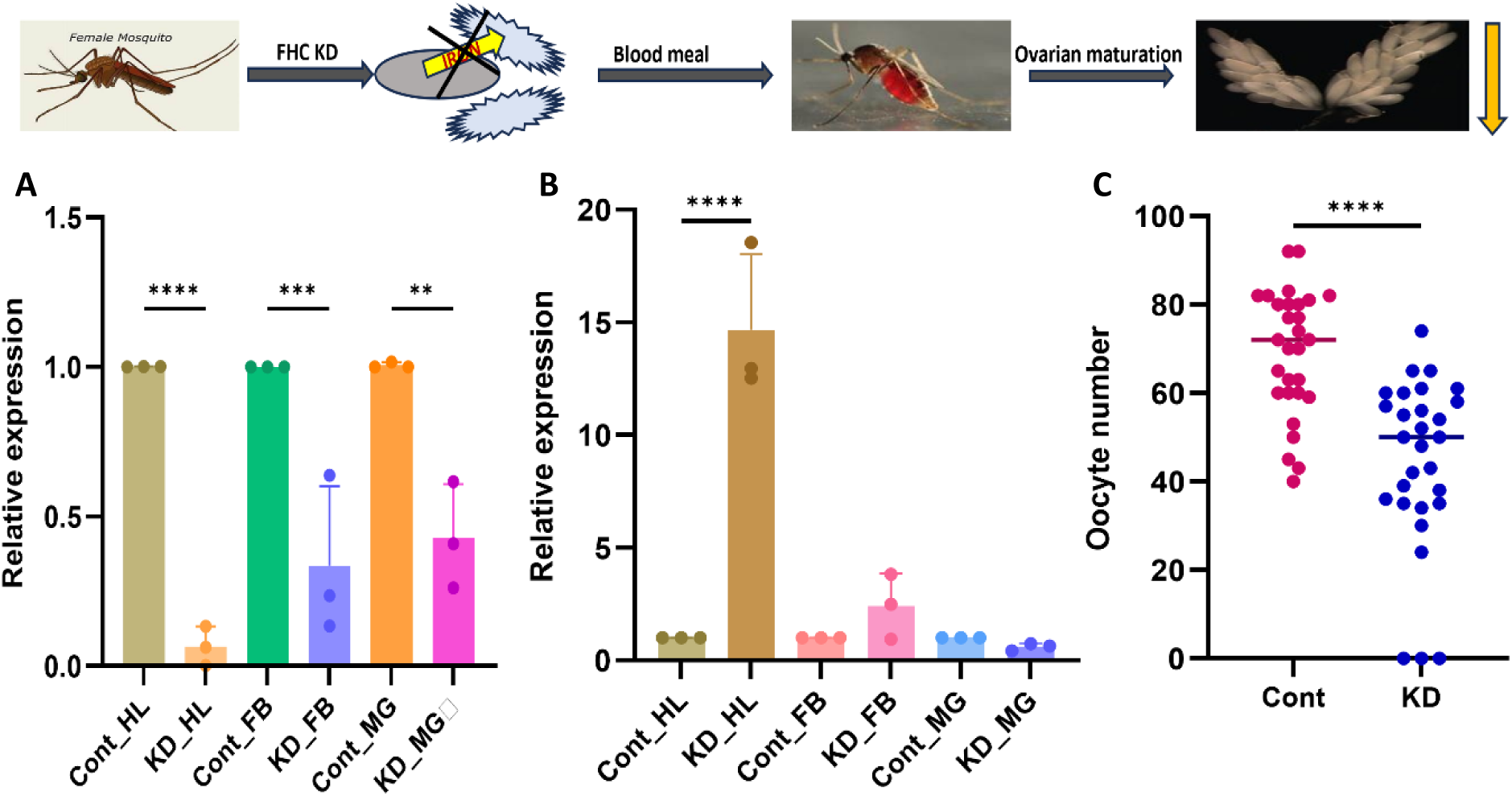
Knockdown effect of *Fer* on oocyte count of mosquito. **(A)** q-PCR based analysis show a major reduction in *Fer* expression in HL (*p* <0.0001), FB (*p* <0.0006) and in MG (*p* <0.0020), after injection of double stranded RNA of *Fer* to 1-2 days old mosquito, taking dsGFP injected as control of same age group (N=3, n= 25). **(B) s**howing increased expression of *Trf* in *Fer* knockdown sample of hemolymph (*p* <0.0001), FB and MG. All three independent biological replicates were statistically analyzed with the help of one-way ANOVA (Tukey’s multiple comparison) test, where asterisks represented *p < 0.05; **p < 0.005; and ***p < 0.0005. (N = number of biological replicates, n number of mosquitoes dissected for sample collection). **(C)** A comparative dot plot graph shows oocyte number reduction in the *dsFer* injected mosquito (*p* <0.0001) as compared to control (*dsGFP* injected) (N=3, n = 10). All three biological replicate data were statistically analyzed by using of Unpaired Two-tailed Student t-test using GraphPad Prism 10, where each dot represents the number of mature ovarian follicles per female mosquito. (N = number of biological replicates, n= number of mosquitoes dissected for sample collection). MG-Midgut, FB-Fat-body, HL- Hemolymph, Cont.- Control, KD- Knockdown.

To further evaluate the possible functional correlation of *Fer* and *Trf* and their role in iron homeostasis regulation, we performed the double knockdown experiment. We mixed the dsRNA of both the *Trf (T)* and *Fer (F)* and injected them, and examined the ovary maturation in the gravid female mosquitoes. In the hemolymph tissue, the expression of the *Fer* and T*rf* genes was significantly reduced to 50% and 70%, respectively (Fig. 3A). Compared to the control mosquito group, *Fer+Trf* KD mosquitoes showed a deleterious impact on their ovarian growth, where (i) 40% of gravid females fail to get complete ovarian follicles maturation (Fig. 3E) (Supplementary Fig. 5A-C); and if they did, (ii) they had 50% reduction in oocytes number (Fig. 3B), and (iii) also a significant size variation in the laid eggs, and the first instar larvae that hatched as well (Fig. 3D, Supplementary Fig. 5E). We noticed control blood-fed mosquito’ hatched larvae have a body size of 751.98 pixels, while the larvae hatched from double knockdown mosquito exhibit a size decrease length by 621.35pixel (Fig. 3C) (Supplementary table 2 & Supplementary Fig. 5D). These data provide direct evidence that a synergistic transcriptional regulation of both Transferrin and Ferritin is crucial to maintain optimal iron homeostasis throughout the vitellogenesis process, and egg development in the gravid female mosquitoes.

**Fig. 3:**
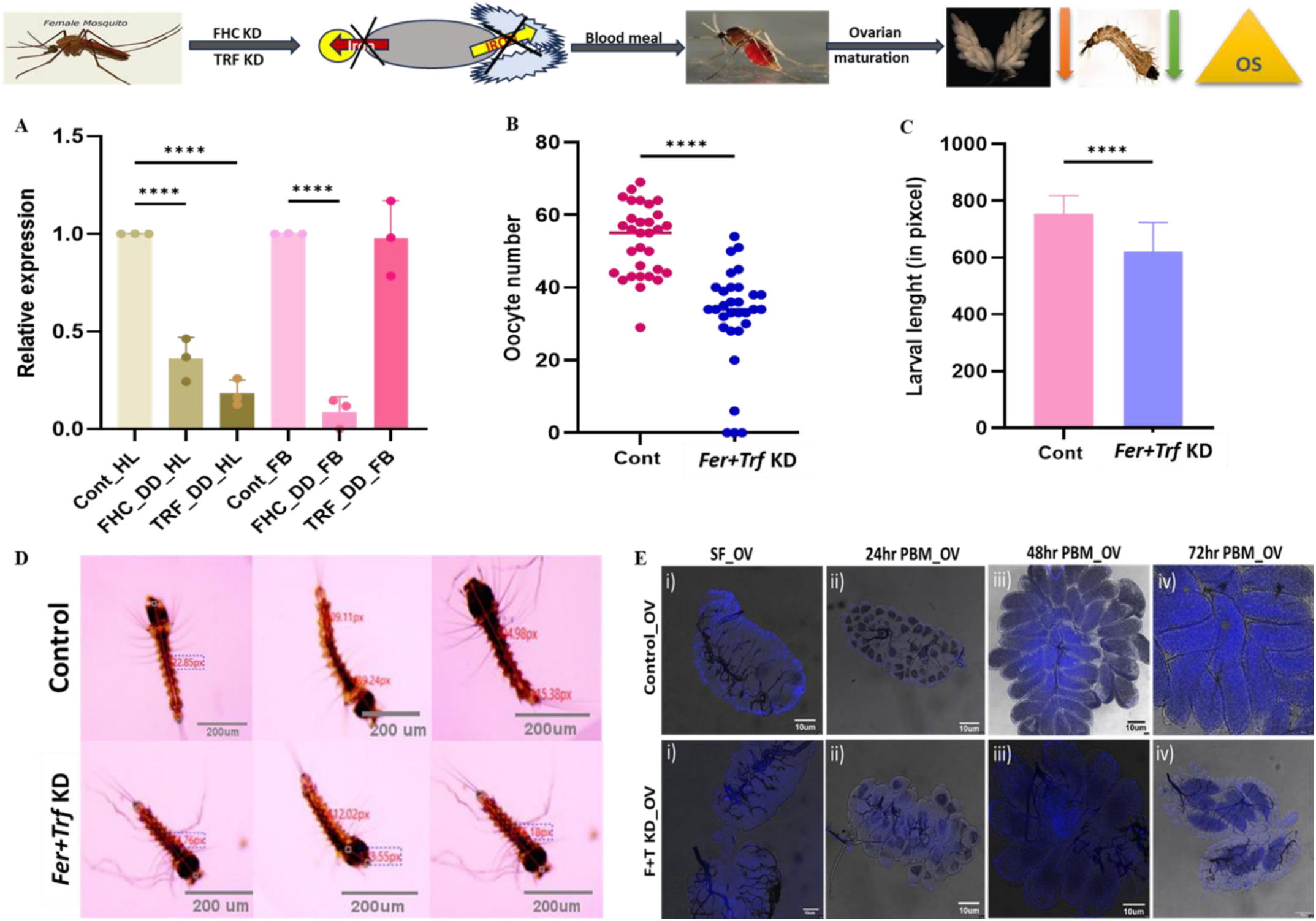
Knockdown effect of both *Fer + Trf* on ovarian development of mosquito: **(A).** q-PCR based analysis show both *Fer* and *Trf* expression reduction in HL and FB (*p* <0.0001) after injection of double-stranded RNA of both *Fer* and *Trf* into 2 days old mosquitoes, taking dsGFP injected as a control of the same age group. (N=3, n= 25). All the three independent biological replicates were statistically analyzed with the help of one-way ANOVA (Tukey’s multiple comparison test) where asterisks represented *p < 0.05; **p < 0.005; and ***p < 0.0005. (N = number of biological replicates, n= number of mosquitoes dissected for sample collection). **(B)** A comparative dot plot graph shows oocyte number reduction in *dsFer+dsTrf* injected mosquito as compared to control (GFP injected). (N=3, n=10). All three biological replicate data were statistically analyzed by using of unpaired two-tailed Student t-test using GraphPad Prism 10 where each dot represents the number of mature ovarian follicles per female mosquito. (N = number of biological replicates, n= number of mosquitoes dissected for sample collection). **(C)** Graph shows difference in larval body size of *dsFer+dsTrf* injected mosquito as compared to control (GFP injected) (N=3, n=7). All three biological replicate data were statistically analyzed by using of unpaired Two-tailed student t-test using GraphPad Prism 10 where asterisks represented *p < 0.05; **p < 0.005; and ***p < 0.0005. (N = number of biological replicates, n= number of mosquitos dissected for sample collection). **(D)** Decreased larval body size was observed in double knockdown mosquito. **(E)** Ovarian development F+T KD mosquitoes (i) Ovaries dissected from sugar fed mosquito control and F+T KD were incubated with DAPI stain for 5 min., washing and visualise (ii) After 24 hr of blood meal ovaries dissected and stain, visualise, (iii) After 48 hr of blood meal ovaries dissected and stain, visualise (iv) After 72 hr of blood meal ovaries dissected and stain, visualise under Leica confocal microscope (Scale bars represent 10 μm). FHC-Ferritin heavy chain, TRF- Transferrin, DD- Double knockdown, HL- Hemolymph, FB- Fat body.

### *Fer+Trf* knockdown modulates gut microbiome and brain responses

Our recent findings highlight that iron-rich blood meal uptake favors rapid establishment of a bidirectional-microbiome-Gut-Brain-Axis (mGBA) communication to maintain optimal organismal physiology till gonotrophic cycle completion in the mosquito *An. culicifacies*(*1*). Because gut bacteria also contribute to iron metabolization, we hypothesized that diminished follicles and poor egg maturation in the *Fer+Trf* KD gravid female mosquitoes could be a result of altered mGBA-associated physiological activities, such as oxidative stress, as proposed in vertebrates, although the underlying mechanism remains unclear(*44*).

To test this, first we evaluated the transcriptional response of nitric oxide synthase (NOS), and stress-responsive enzyme i.e. superoxide dismutase (SOD) in the brain and gut, respectively, of control as well as *Fer+Trf* KD adult female mosquitoes. Interestingly, we observed at least 2.5-fold enrichment in the NOS expression (Fig. 4B), and ROS-induced fluorescence, as detected by DCFDA dye, in the brain of the non-blood-fed *Fer+Trf* KD mosquito (Fig. 4A). This indicated *Fer+Trf* expression dysregulation may likely interfere with the NOS-mediated neuro-signaling in naïve mosquitoes. And when given blood-meal, except for early induction (6hr) in the control, the expression of NOS remains static till 24hrs in the brain of both groups of mosquitoes (Supplementary Fig. 7 B-C), indicating KD *Fer+Trf* has a direct influence on the brain response. Surprisingly, noticing a 2.0-fold enrichment of SOD in the gut of the *Fer+Trf* KD blood-fed mosquitoes (Fig. 4F), further indicated that iron-homeostasis disruption may lead to increased free radicles *via* Fenton reactions in the gut. Finally, a notable enrichment of 5HT, a receptor for the neurotransmitter serotonin (Fig. 4D), which performs various brain functions including learning, memory and feeding behaviour, together confers that impairing iron-metabolism not only limits the iron availability to brain, may also alter the expression oxidative stress responsive effectors influencing GBA communication(*45*).

**Fig. 4:**
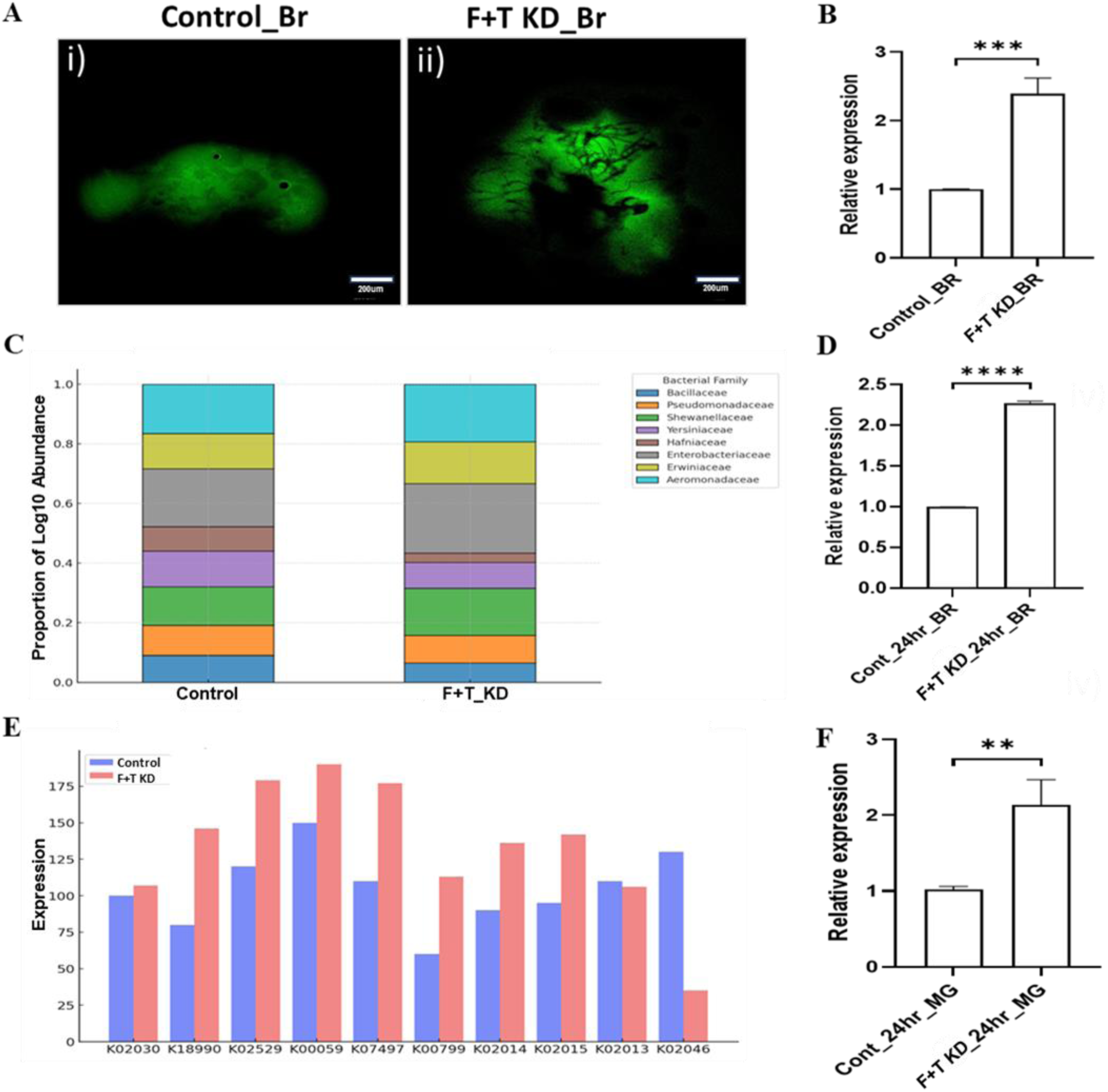
Iron-homeostasis disruption modulates gut microbiome and brain responses: **(A)** The mosquito brain was incubated with CM-H_2_DCFDA (2 µM) for 20 min, washed, and visualized under a confocal microscope. Scale bar– 200 µm. (i) ROS staining of control sugar-fed brain with dsGFP injected, (ii) Sugar-fed F+T KD brain showing different patterns of ROS production. Scale bar – 200 µm. (also see supplemental Table 5); **(B)** Real-time expression analysis of NOS in double KD brain samples show increased expression of NOS (*p*<0.0001) after injection, taking dsGFP injected as control of the same age group. (N=3, n= 25). **(C)** Taxonomic Abundance in Control vs F+T KD Samples (Kraken2 Analysis): This stacked bar chart displays the taxonomic abundance of bacterial families in Control vs F+T KD (D0) samples, analyzed using Kraken2. The Y-axis represents the proportion of log10 abundance, with different colours indicating distinct bacterial families. This comparison highlights shifts in microbial composition due to F+T KD, revealing potential microbial community alterations. **(D)** Real-time expression analysis of 5HTR in 24hr blood fed double KD brain samples show increased expression of 5HTR (*p*<0.0001) after injection, taking dsGFP injected as control of same age group. (N=3, n= 25). All three biological replicate data were statistically analyzed by using of unpaired two-tailed Student t-test using GraphPad Prism 10 where asterisks represented *p < 0.05; **p < 0.005; and ***p < 0.0005. (N = number of biological replicates, n= number of mosquitoes dissected for sample collection). **(E)** Gene Expression Levels in Control vs F+T KD Samples: This bar chart represents the variation in predicted metabolic pathway linked gene expression levels between Control and F+T KD samples. Gene expression was measured based on read-level taxonomic profiling, where processed paired-end reads were aligned to the NCBI 2019 Genome Build using KMA. The red bars represent F+T KD samples, while the blue bars represent Control samples. Differences in expression suggest significant regulation of specific genes following F+T KD. **(F)** Real time expression analysis of SOD in 24hr blood fed double KD midgut samples show increased expression of SOD (P<0.0047) after injection, taking dsGFP injected as control of same age group. (N=3, n= 25). All three biological replicate data were statistically analyzed by using of unpaired two-tailed Student t-test using GraphPad Prism 10 where asterisks represented *p < 0.05; **p < 0.005; and ***p < 0.0005. (N = number of biological replicates, n= number of mosquitoes dissected for sample collection) and a * mark was drawn automatically by the software by analyzing their significance itself. (N = number of biological replicates, n= number of mosquitoes dissected for sample collection).

Further, observation of alteration in the total gut-bacterial population of the blood-fed *Fer+Trf* KD mosquitoes (Supplementary Fig.7A), prompted us to evaluate whether iron homeostasis disruption also interferes with iron metabolization-associated bacterial abundance. To examine the microbial properties and functional diversity of the gut-bacterial population, we carried out WGS-based comparative metagenomic analysis (Fig. 4C). Interestingly, a close review endorsed that *Fer+Trf* KD has a greater influence on specific bacterial community, probably linked to iron metabolism, such as *Aeromonadaceae* (genus *Aeromonas)*, *Enterobacterales (Enterobacter*), and their family members. Further, we also observed a restricted growth of certain microbial species, including *Aeromonas hydrophila, Enterobacter hormaechei*, and *Klebsiella aerogenes*, while favoring the growth of others, such as *Klebsiella pneumoniae* and *Enterobacter asburiae* (Supplementary Fig. 6A-D). A comparative pathway prediction analysis highlighting relative metabolic changes (Fig. 4E) and associated enzyme enrichment, corroborates that the iron-homeostasis disruption modulates specific functional properties of bacteria, likely iron metabolism and oxidative responses (Supplementary Fig. 6E).

### Mosquito *Hepcidin*

Mammalian systemic iron homeostasis is mainly regulated by the hormone hepcidin(*46*), which is dominantly synthesised by the liver(*47*). But in case of insects, it is proposed that they do not code for similar proteins, and hence may likely have different regulation mechanisms(*48*). However, while re-analysing and updating our pre-existing hemocyte RNA-seq data (NCBI accession numbers SRR12031469;(*3*)We unexpectedly discovered at least three independent transcripts encoding bony fish homologs of putative Hepcidin, Ceruloplasmin, and Transferrin-like proteins (Supplemental Table 4). A detailed homology search failed to show similarity against any insect/mosquito species database (Supplemental Fig. 8B, D). While an independent analysis is ongoing to unravel the possible evolutionary relationship and acquisition mechanism (Rani et al., Unpublished), here, we evaluated whether the identified novel hepcidin has any functional correlation in iron homeostasis regulation.

A comprehensive *in-silico* analysis reveals that the identified putative 222bp long encodes a partial protein (59 AA) and lacks a cysteine-rich fragment (Fig. 5 A, B, C, Supplemental Fig. 8A). Furthermore, a low expression was only detectable by RT-PCR in the fat body, which was slightly enhanced in *Fer+Trf* KD mosquitoes, when evaluated by Real-Time PCR assay (Supplementary Fig. C, E). Thus, a detectable expression in the fat body (equivalent to the mammalian liver), suggests that the identified Bony-fish homolog (*AcHep*) may have some role in the systemic iron homeostasis regulation in this mosquito species, though it is yet to be explored.

**Fig. 5.**
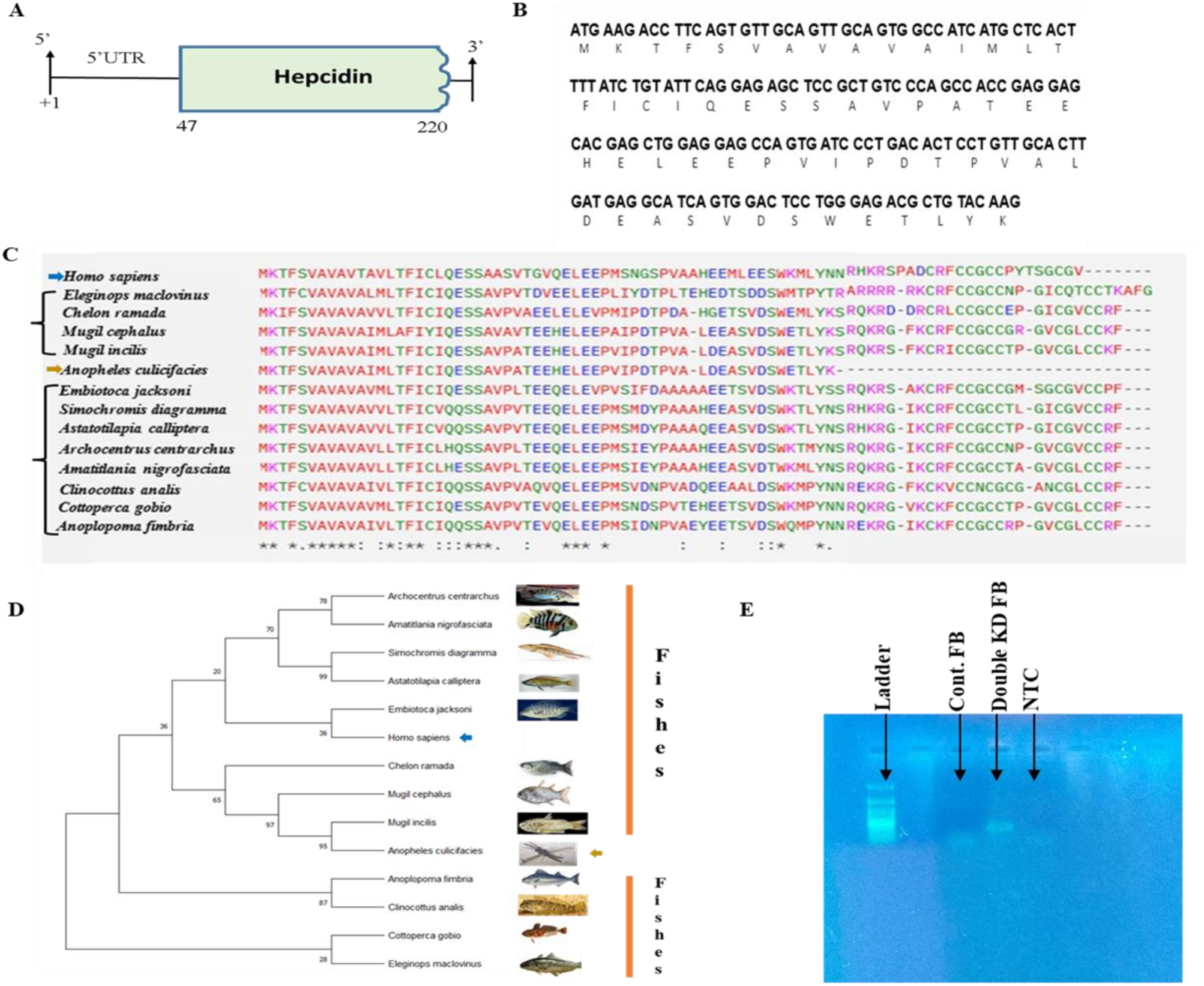
Sequence analysis of *An. culicifacies* transcript *Hepcidin* and its transcriptional response. **(A-B)** Genomic overview and molecular features of identified partial cDNA sequence of mosquito Hepcidin (green region highlights coding sequences); The gene contains 222bp nucleotide, encoding 59 AA long peptides. **(C)** Multiple sequence alignment performed by Clustal Omega showing gaps and dissimilar amino acids in homologous species. The multiple sequence alignment of Hepcidin shows homology with different vertebrate species including *Homo sapiens*, rather than insect. Here, the asterisk ‘*’ shows the conserved amino acid, ‘.’ Showing some conservation, **‘-’** showing gap means no amino acid, whereas ‘:’ showing amino acid of one group. Control. **(D)** The phylogenetic tree of Hepcidin was obtained using by Molecular Evolutionary Genetics Analysis (MEGA) tool. The maximum likelihood method was applied along with 1000 bootstrap iterations and the Jones-Taylor-Thornton model of amino acid substitutions. **(E)** Gel picture showing a light band in the fat-body of *Fer+Trf* KD mosquitoes, whereas no band appears in control and No-Template Control (NTC)

## Discussion

Here, we investigated the role of transferrin and ferritin in the regulation of systemic iron homeostasis and reproductive physiology of the mosquito *An. culicifacies*, an important rural malarial vector in India. We report that a coordinated transcriptional regulation of these iron-binding proteins are crucial for iron mobilization and reproductive success and when dysregulated, it impairs ovarian follicle maturation and egg development, likely by altering the microbiome-gut-brain-axis communication in the gravid female mosquitoes. Furthermore, our transcriptomic analysis identified an unreported putative hepcidin-like transcript in *An. culicifacies*, suggesting a potentially novel factor in mosquito iron regulation. Although its functional role has not been determined, its presence highlights the possibility of an uncharacterized mechanism in mosquito iron homeostasis.

In mammals, iron is primarily bound to haemoglobin and myoglobin where it plays a vital role in oxygen transport and storage. Dysregulation in iron homeostasis results in various pathological conditions, including metabolic syndrome, sepsis, iron deficiency, and inflammation(*49*). Among several reported mammalian iron-mobilizing factors, ferritin and transferrin significantly contribute to systemic iron homeostasis(*50*, *51*).

Hematophagous insects, including mosquitoes, consume a large amount of blood for their reproductive success(*52*). And these insects have developed multiple mechanisms to combat the large concentration of heme present in the ingested blood meal. Heme-derived iron majorly contributes to the eggs, where ferritin likely mediates its storage and transport between haemolymph to eggs(*53*). In the cattle tick, *Boophilus microplus*, heme metabolism (a prosthetic group containing iron) has been described(*54*) and a secreted protein, Ferritin2, has been suggested to participate in non-heme iron metabolism in the tick *Ixodes ricinus*(*12*). In *Aedes aegypti,* iron transporters ZIP13 seems to play an important role in the gut-iron mobilization, but have no effect on ovary maturation, pointing to an alternative mechanism of heme-derived iron deployment that may exist in mosquito(*55*).

After the blood meal acquisition, mosquitoes experience a physiological shift that activate multiple organs to synchronize digestion and reproduction(*56*). Recently, we have demonstrated the importance of microbiome gut-brain-axis orchestrating physiological homeostasis during the blood meal digestion and vitellogenesis process in the mosquito *Anopheles culicifacies*(*1*). Gut-inhabiting microbiota regulates many physiological processes in the host but their role in gut iron metabolism remains unclear(*26*). Once transported to haemolymph, iron derived from blood meal is either stored in the fat body or mobilized to facilitate ovarian development. Considering these findings, we envisioned that any dysregulation in the iron mobilization may have a direct impact on ovarian growth and development.

To test this proposition, we are currently evaluating the functional correlation of two well-known iron-storage and mobilizing factors, ferritin and transferrin, in the mosquito *An. culicifacies*. We have shown that iron transporter *transferrin* significantly contributes to iron mobilization during the ovarian follicle maturation(*3*). While investigating the role of ferritin, like transferrin, we noticed that a high expression of ferritin occurs in the eggs than in other aquatic stages, and adult females than males, which together underscore their evolutionary adaptation of efficient iron mobilization and utilization in hematophagous female mosquitoes. Tissue-specific expression modulation after silencing either of transferrin or ferritin indicated that a multi-organ level coordination across the fat-body, hemolymph, and ovaries is important to maintain a basal level of iron homeostasis. Further correlating to transferrin, data on spatial/temporal expression modulation in response to blood meal and reduced oocyte development after silencing of ferritin, ensured that a synergetic transcription of ferritin/transferrin could be crucial for systemic iron-homeostasis regulation and ovary maturation in gravid female mosquitoes.

To validate the hypothesis, we disrupted this synergistic regulation by silencing both transferrin and ferritin and assessed its effect on reproductive outcome in the gravid females. Astonishingly, we observed a dramatic impact on the developmental failure of ovarian follicles in at least 40% of females, and if they delivered, they had reduced egg number and size variation in the hatched first instar larvae. These findings support the conceptual idea that systemic iron homeostasis regulation in hematophagous insects may be more akin to that of mammals, especially iron-deficiency and pregnancy in humans(*57*).

Because iron is crucial for both maternal and foetal health during pregnancy in mammals, its deficiency not only causes increased maternal illness risks but also adversely affects reproductive outcome, including intrauterine growth restriction, prematurity, poor foetal development with low birth weight, and neuro-disorders(*58*). While estimation of serum ferritin and transferrin saturation levels serves as a key marker for Iron deficiency assessment in pregnancy, only limited studies in mouse/animal models correlate their impact of iron deficiency and placental growth or foetal development. The cross-tissue iron homeostasis regulation mechanism and its physiological relevance remain questionable(*59*).

Pregnancy is susceptible to oxidative stress; any imbalances in transferrin–ferritin regulation and hemolysis may lead to oxidative stress and damage of cellular components, particularly placental membranes, promoting ferroptosis and miscarriage in women(*60*, *61*). Emerging literature further suggests that gut microbiota significantly contributes to host iron absorption and homeostasis, but understanding the nature of interaction and their cross-talk during iron deficiency-related anaemia remains a major challenge(*62*).

The intake of iron, deficiency or excess, in the host influences the composition of microbiota. Based on this, we hypothesized that alteration in iron-mobilization may alter the gut-microbiome population and, therefore, neuro-communication, the ultimate cause of increased oxidative stress and iron toxicity affecting the mosquito reproduction. A comparative WGS-metagenomic analysis not only showed a greater influence on specific bacterial communities, namely *Aeromonadaceae* (genus *Aeromonas)*, *Enterobacterales (Enterobacter*) but also a remarkable change in gut-microbiota-associated metabolic activities linked to iron metabolism and oxidative responses. As expected, we noticed an altered ROS signal and titer of nitrite in the brain during early hours of blood-feeding, and enhanced SOD expression level in the gut in the *Fer+Trf* KD mosquitoes.

Studies in mammals highlight that iron deficiency significantly alters the 5-HT level, and a disruption in the serotonin level may trigger multiple neurotransmission activities, including neuro-behavioral manifestations, neuropsychological disorders, and depression(*63*, *64*). Previously, 5-HT receptors have been implicated in the regulation of male mosquitoes’ auditory behavioural responses(*65*), whereas in female mosquitoes, it has been suggested to influence salivation and gut-microbial interaction, parasite transmission, and ovarian development(*66–68*). Although in mosquitoes, blood-feeding significantly modulates neurotransmitter (NTs) level and their receptor expression in the gut and brain of mosquitoes, their physiological relevance to iron metabolism remains uncertain(*1*). Therefore, we tested whether iron homeostasis dysregulation affects the 5-HT expression in the brain and gut of the mosquitoes. Interestingly, we noticed blood-feeding significantly upregulates, while silencing of *Fer+Trf* downregulates the 5HT expression, in the brain of adult female mosquitoes. We have previously observed that blood-feeding to the aseptic (antibiotic-treated) mosquitoes results in a decrease in expression of 5HT in the brain and gut of adult female mosquitoes(*1*). Our findings not only corroborate the 5HT role in neuro-regulation, as in mammals, but also highlight that a systemic iron-homeostasis dysregulation may significantly affect Microbiome-Gut-Brain-Axis communication, and hence reproductive physiology, in the gravid female mosquitoes.

Interestingly, while the insect genome is supposed not to encode for hepicidin protein-a peptide hormone essential for iron regulation in mammals, we discovered a bony fish homolog hepicidin-like transcript in the hemocyte RNAseq data of *An. culicifacies*(*69*, *70*). Although hepcidin expression was undetected in the midgut and ovaries, its expression was detectable only in the fat body of the *Fer+Trf* silenced mosquitoes. Further, functional analysis is in progress to predict and evaluate their possible evolutionary significance and possible link with iron homeostasis in the mosquito *An. culicifacies*.

## Supporting information

Supplemental data sheet

## Authors contribution statement

PY, SK, ST and RD conceived and designed the experiments; PY, JR, VS, PR, VS, NS, GS, TS, GT, STY helped to perform experiments, drafting and editing MS; PY, JR, SK, ST, RD Contributed reagents/materials/analysis tools, helped in writing, editing, reviewing and finalizing MS.

## Acknowledgement

We would like to thank the ICMR-National Institute of Malaria Research insectary staff members for mosquito rearing. We thank Kunwarjeet Singh for their technical assistance in the laboratory. We also thank MOLSYS SCIENTIFIC for WGS shotgun sequencing.

## Data Submission Detail

The metagenomic sequence data have been submitted to the NCBI SRA database (SAMN47311708)

