## Supplemental data sheet for "Iron homeostasis and reproduction: Unveiling the Microbiome-Gut-Brain-Axis connection in the mosquito *Anopheles culicifacies*"

**Table 1: List of primer and their sequences.**

| S.No. | Name of the gene | Left primer | Right primer |
|---|---|---|---|
| 1. | Ferritin Heavy chain (Fer) | GAAGCTGATCGAATATGCTC | GGTAGTCGACCAGATGGTAA |
| 2. | Transferrin (Trf) | AGTTTGAGATAGGCAGCGA | CATCCACCGCATTCATGAA |
| 3. | Fer_DSR | TAATACGACTCACTATAGGGCCTACTTTGCCCAGTACAAG | TAATACGACTCACTATAGGGTAATCGTTGTGATCCTCCTC |
| 4. | Transferrin_DSR | TAATACGACTCACTATAGGGCAGGACCATGACAAGTTCG | TAATACGACTCACTATAGGGCTTGCGGCGCTTCTTTAT |
| 5. | NOS | ATGAGGACCAACTATCGGG | GCCTTGGTGACAATGCTC |
| 6. | SOD3 | TAGAAGGTTTACGACCAGGA | ATGATGTTACCGAGATCACC |
| 7. | Hemeoxygenase 1 | GCGCAAAATAGTAGACGAGT | AACTGCGTTAAAACGATACC |
| 8. | 5HT | ATGATCTCGCGTAACTCCTC | ATCGGATTGACCAGACTGC |
| 9. | 16S | TTGGAGAGTTTGATCCTGGCTC | ACGTCATCCCCACCTTCCTC |
| 10. | Ac_ACT | GCGGTATACTGACACTCAA | CAAACATGATCTGTGTCATC |
| 11. | Ac_RSP7 | ATCGCTATGGTGTTCGGTTC | TTGTTGAACTCGACCTCACG |
| 12. | Ac_hep | CAAGAATCACCAGGAGATCC | TTGTACAGCGTCTCCCAG |

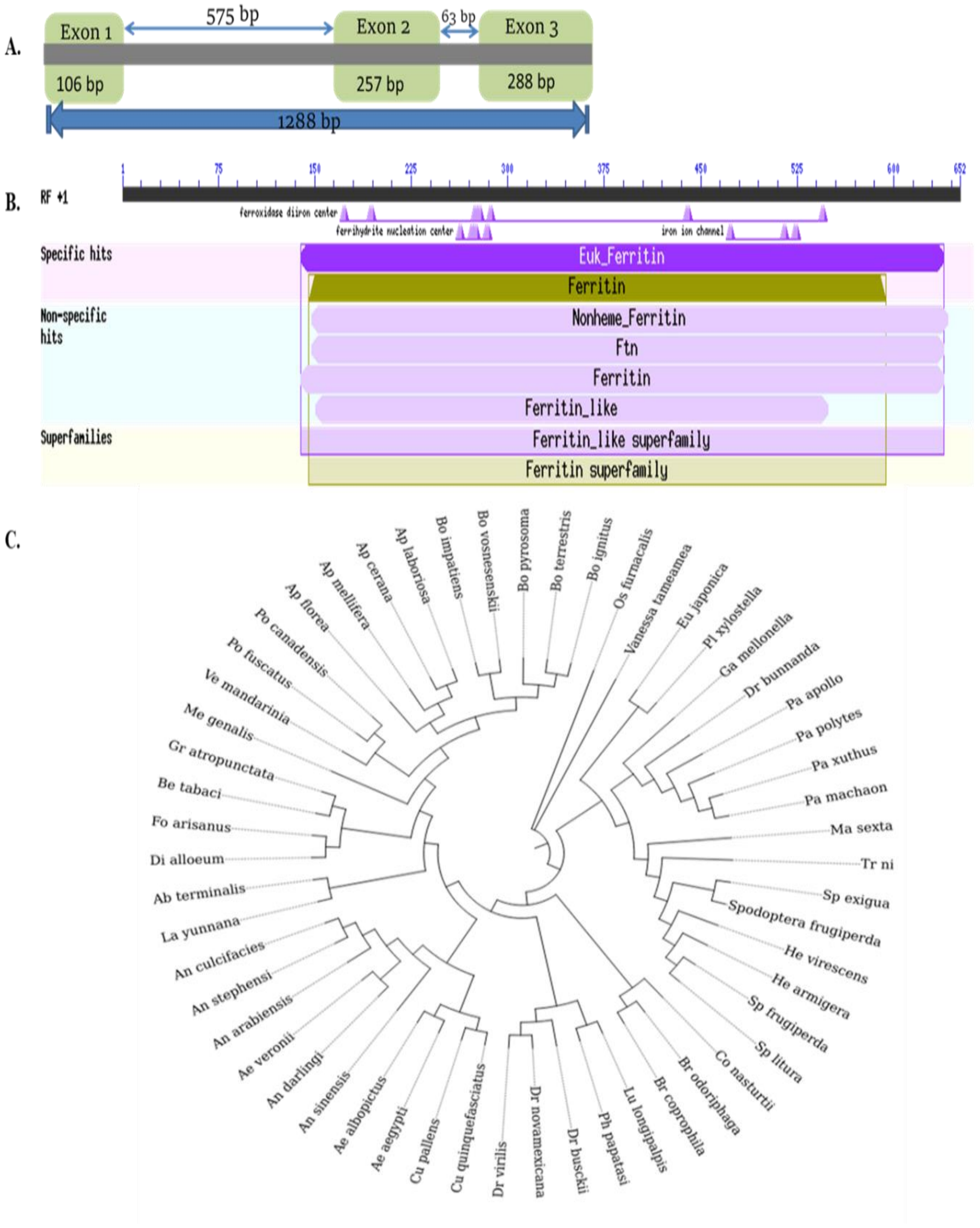


**Fig. 1: In-silico analysis of Ferritin *(Fer)*.** **(A)** Genome organization of Ferritin (*Fer*) in *An. culicifacies* showing three exons and 2 introns are present in the genomic sequence of the transcript. **(B)** Domain annotation of *An. culicifacies* Ferritin *(Fer)*. **(C)** Phylogenetic analysis of selected transcripts was done using the maximum likelihood algorithm with 1000 times bootstraps in MEGAX software

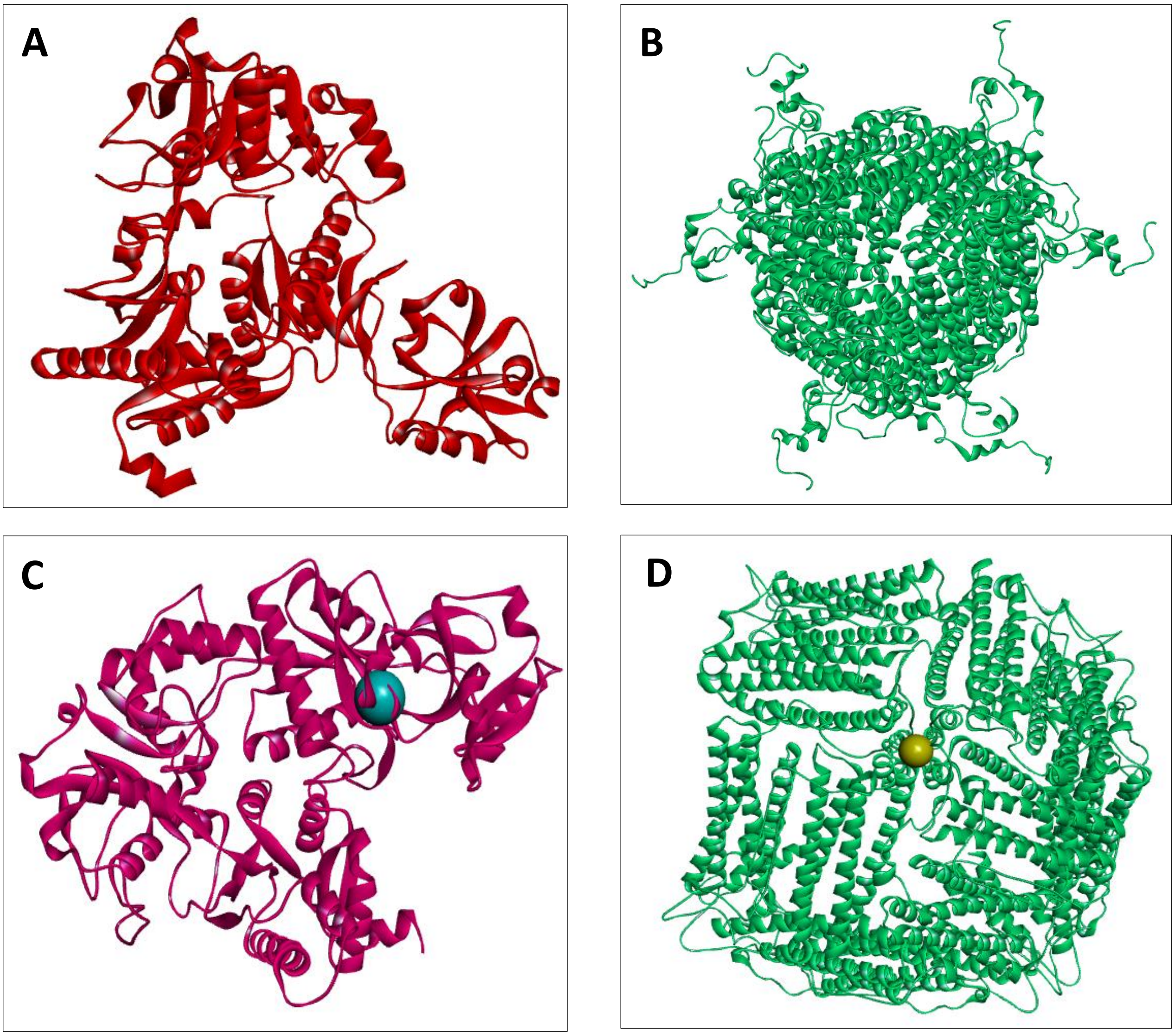


**Fig. 2: Sequence analysis and Modelling of Ferritin and Transferrin. (A)** Final 3D structure of Transferrin was constructed using GalaxyHomomer Web Server (http://galaxy.seoklab.org/homomer). **(B)** Ferritin 3D model of ferritin was constructed using Modeller version 10.5. Models obtained were further refined and validated with PROCHECK and ModRefiner respectively. After the 3D structures were finalized, active sites of Ferritin and Transferrin were predicted using CASTp (Computed Atlas of Surface Topography of proteins). **(C)** Transferrin bind with blue round ball indicates ferric ($Fe^{3+}$) ion while **(D)** The yellow round ball is ferrous ($Fe^{2+}$) ion bound to Ferritin. Docking of $Fe^{2+}$(ferrous ion) and $Fe^{3+}$(ferric ion) done using molecular docking program AutoDock vina 1.5.7. Grid around the packets was created and adjusted according to the number of points in X, Y, Z-axis so that the entire active site of the proteins is covered. The default value is 0.375 Å between grid points, which is about a quarter of the length of a carbon-carbon single bond. For studying the interaction between the proteins and the ions PyMol and Discovery Studio v. 24.1.0 was used.

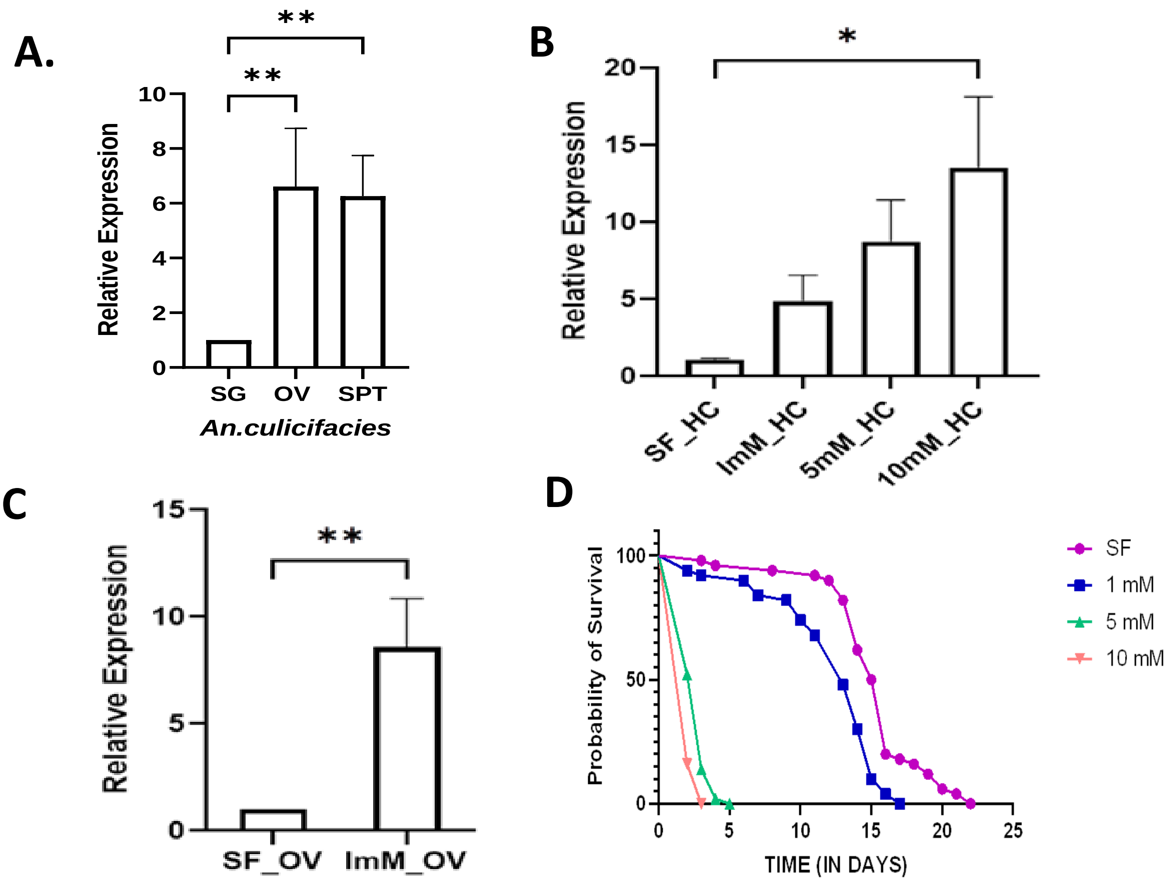


**Fig. 3: Exogenous iron supplementation affects the ferritin expression and mosquito survival (A).** High transcript level in female reproductive organs, considering the salivary gland as a control for all test samples. (N=3, n=25); **(B)** Relative Ferritin *(Fer)* expression after exogenous supplementation of iron in different concentrations (1mM, 5mM, and 10mM) in hemolymph ($p<0.0426$); and in **(C)** the ovary ($p<0.0004$) shows increased expression of Ferritin *(Fer)* with increasing concentration of iron. Here, sugar-fed was considered as control for all test samples. (N=3, n=25). All the three independent biological replicates were statistically analyzed with the help of one-way ANOVA, where asterisks represented *$p < 0.05$; **$p < 0.005$; and ***$p < 0.0005$. (N = number of biological replicates, n= number of mosquitoes dissected for sample collection). **(D)** Survival graph shows high iron concentration affects the survival of mosquito by increasing toxicity. (N=3, n =25). All three independent biological replicates were statistically analyzed by using GraphPad Prism 10 software. (N = number of biological replicates, n number of mosquitoes dissected for sample collection).

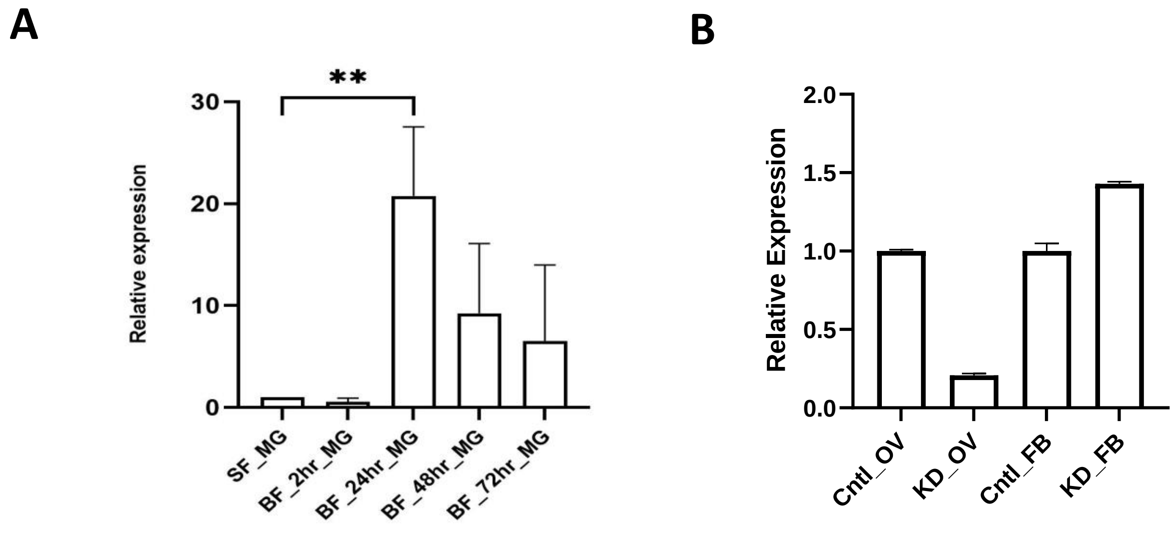


**Fig. 4: (A)** Heme oxygenase (HO) expression: Highest expression of heme-oxygenase at 24hr post blood meal in midgut (MG) of mosquito, where sugar-fed midgut was taken as control. **(B)** The graph shows Ferritin expression in Transferrin knockdown samples (N=3, n=25). All three independent biological replicates were statistically analyzed with the help of one-way ANOVA where asterisks represented *$p < 0.05$; **$p < 0.005$; and ***$p < 0.0005$. (N = number of biological replicates, n= number of mosquitoes dissected for sample collection).

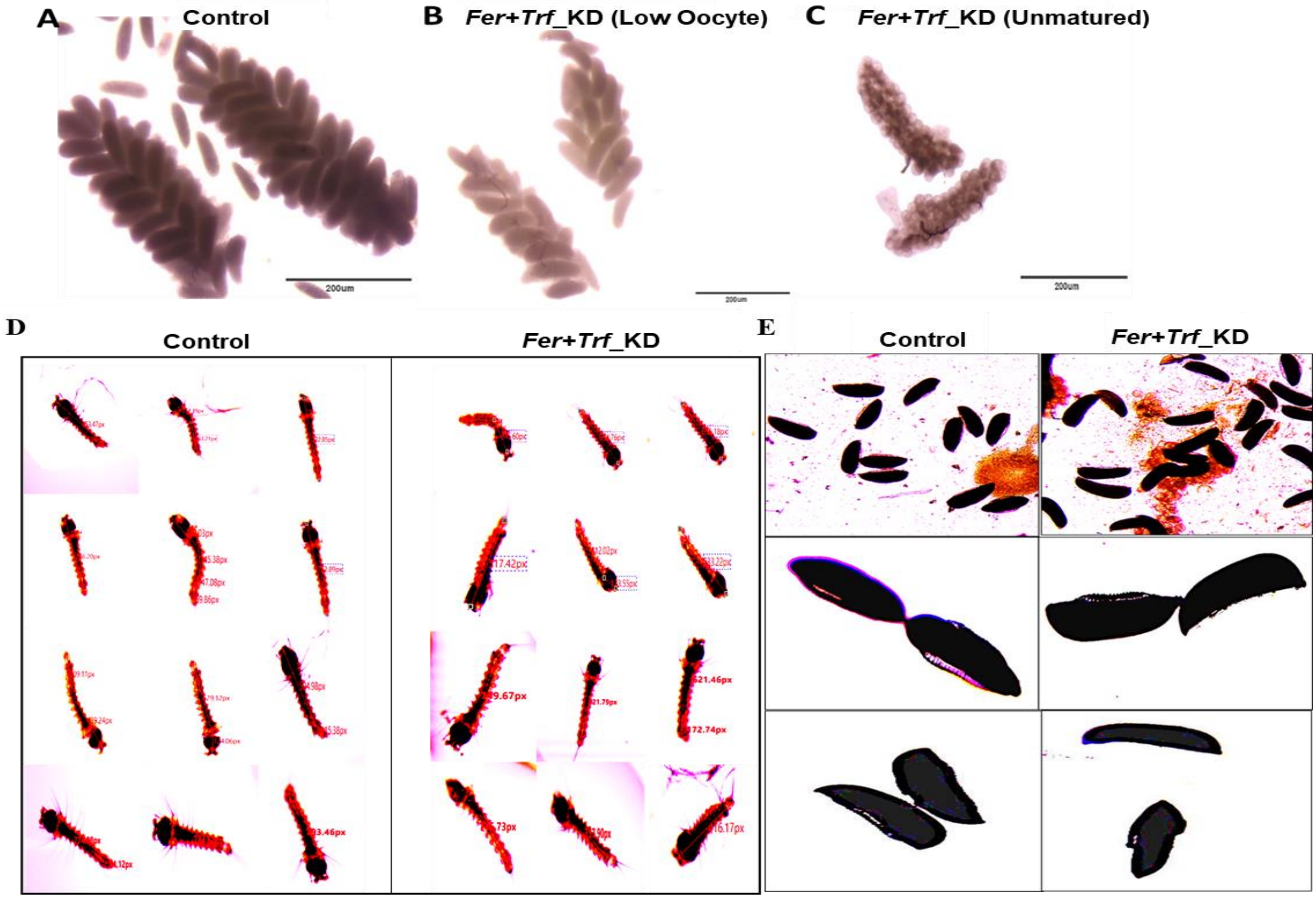


**Fig. 5: Microscopic ovarian assessment and larval body size of the mosquito.** After 60 hours of blood meal, mosquito of both control and knockdown were dissected, and phase-contrast microscopic photographs of the ovaries were taken at 4X magnification. **(A)** Fully matured ovary of a control mosquito. **(B)** Low number of oocyte count in the double knockdown mosquito. **(C)** Immature ovary in a double knockdown mosquito after 60 hours of blood meal. Scale bar added by using software IMAGEJ. **(D)** Decreased larval body size was observed in the double knockdown mosquito. (N=3, n=7). Image was captured by a camera fitted microscope, and with Magvision software at 4X and length measurement was also done by using Magvision measurements. **(E)** Size difference in a laid egg of the control and Fer+Trf KD mosquito.

**Table 2:** Showing different larval lengths in control as well as double knockdown samples.

| S.No. | Length of control larvae (in pixcel) | Length of *Fer+Trf* kd larvae (in pixcel) |
|---|---|---|
| 1. | 653.47 | 569.62 |
| 2. | 695.6 | 534.76 |
| 3. | 722.85 | 535.18 |
| 4. | 650.35 | 517.42 |
| 5. | 752.89 | 525.57 |
| 6. | 748.35 | 416.17 |
| 7. | 753.18 | 533.19 |
| 8. | 845.17 | 655.13 |
| 9. | 674.59 | 481.8 |
| 10. | 726.42 | 642.32 |
| 11. | 742.44 | 665.46 |
| 12. | 710.82 | 649.98 |
| 13. | 844.76 | 706.82 |
| 14. | 818.37 | 757.9 |
| 15. | 838.43 | 746.73 |
| 16. | 726.42 | 642.32 |
| 17. | 730.87 | 683.01 |
| 18. | 737.43 | 651.3 |
| 19. | 828.11 | 749.67 |
| 20. | 839.1 | 762.76 |

**Table 3: Showing mosquito Hepcidin identity with other vertebrate hepcidin.**

| **Species** | **No. of Amino_acids** | **Percent identity** | **Coverage** | **Accession No** |
|---|---|---|---|---|
| ***Mugil incilis*** | 85 | 88 | 79 | UXV25347.1 |
| ***Mugil cephalus*** | 85 | 89.66 | 79 | XP_047454919.1 |
| ***Embiotoca jacksoni*** | 86 | 77.5 | 78 | XP_068996370.1 |
| ***Archocentrus centrarchus*** | 86 | 68.9 | 78 | XP_030605780.1 |
| ***Amatitlania nigrofasciata*** | 86 | 68.9 | 78 | AHF46363.1 |
| ***Cottoperca gobio*** | 73 | 72.4 | 78 | XP_029306823.1 |
| ***Anoplopoma fimbria*** | 86 | 72.4 | 78 | XP_054483003.1 |
| ***Chelon ramada*** | 85 | 77.5 | 79 | QBO59819.1 |
| ***Eleginops maclovinus*** | 89 | 70.6 | 78 | ABY84826.1 |
| ***Clinocottus analis*** | 86 | 65.52 | 78 | XP_068425196.1 |
| ***Simochromis diagramma*** | 86 | 74.14 | 78 | XP_039868628.1 |
| ***Astatotilapia calliptera*** | 86 | 74.14 | 78 | XP_026040957.1 |
| *Anopheles culicifacies* | 73 | 100 | 100 | |
| ***Homo sapiens*** | 84 | 58.6 | 78 | AAT74401.1 |

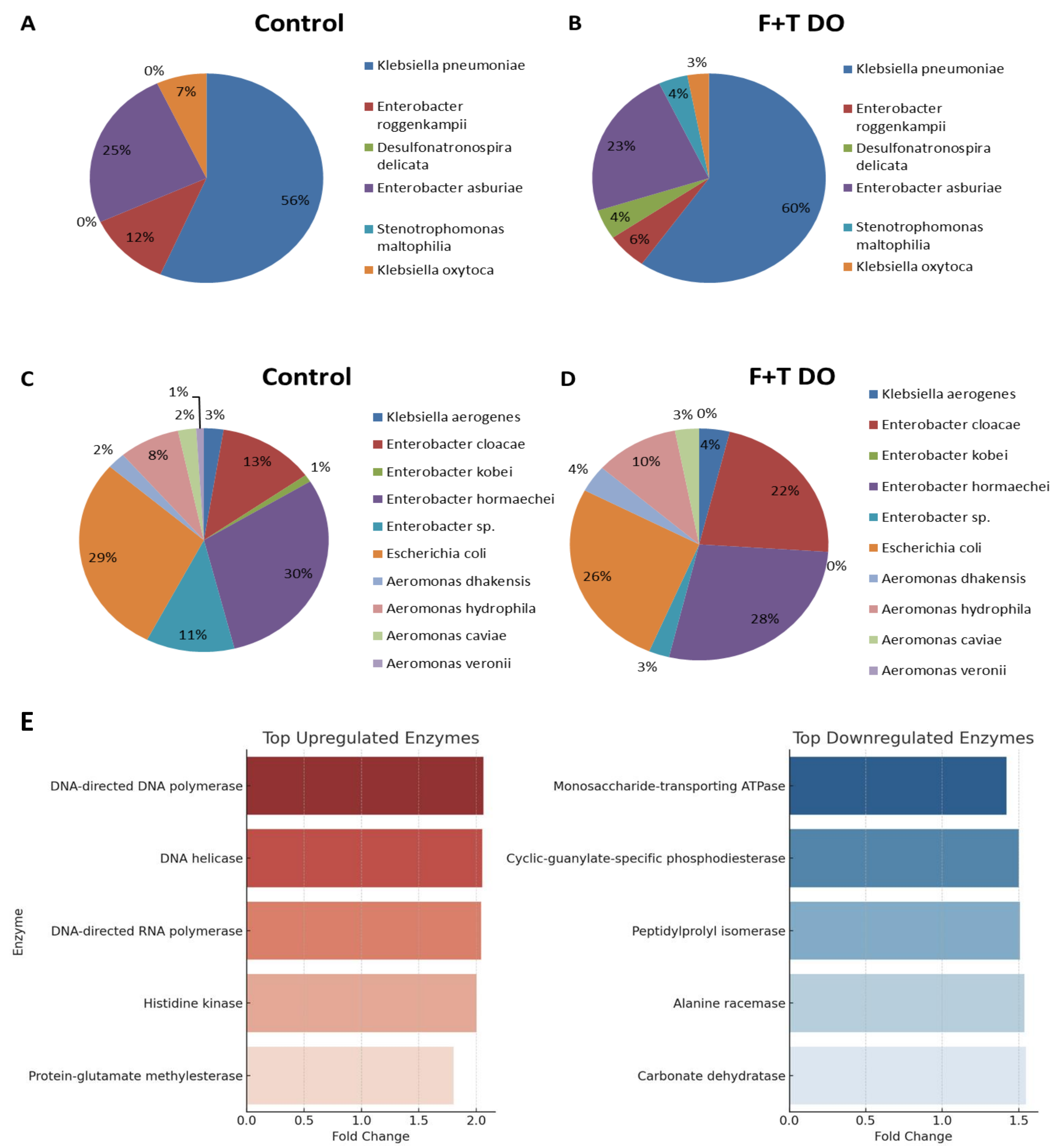


**Fig. 6: Bacterial abundance difference in Control and F+T knockdown samples.** Comparative pie chart showing the difference in gut bacterial abundance increased after F+T Knockdown. **(A)** Bacterial abundance in Control **(B)** F+T Knockdown. Comparative pie chart showing the difference in gut bacterial abundance decreased after F+T Knockdown. **(C)** Bacterial abundance in control **(D),** F+T Knockdown. **(E)** Bakta-Annotated Enzyme Fold Regulation Based on Abundance: Double heatmap showcases the top upregulated (left, red) and downregulated (right, blue) enzymes, annotated using Bakta, based on fold-change regulation in response to F+T knockdown. Upregulated enzymes (left) show increased abundance in F+T DO samples compared to Control. Downregulated enzymes (right) exhibit reduced abundance after ferritin knockdown.

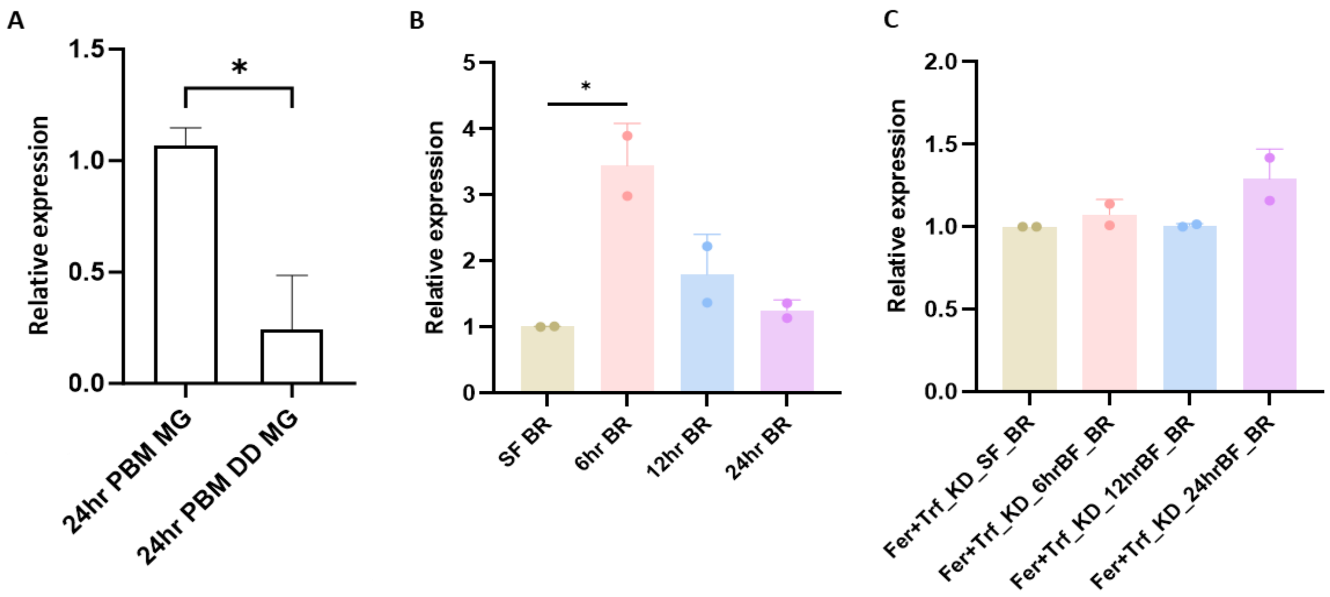


**Fig. 7: *Fer+Trf* KD effect on bacteria and oxidative stress: (A)** bacterial population determined through 16S transcript expression: Real time expression analysis of 16S after 24hr of blood feeding midgut from both control and *Fer+Trf* KD and found a minor reduction in bacterial population (*p* <0.0078) (N=3, n=25). All three independent biological replicates were statistically analyzed with the help of one-way ANOVA (Dunnett's multiple comparison test) where asterisks represented *p < 0.05; **p < 0.005; and ***p < 0.0005. (N = number of biological replicates, n= number of mosquitoes dissected for sample collection). **(B)** NOS expression analysis in SF brain as a control and different time point like 6hr having higher transcript level (*p* <0.0182), 12hr, 24hr after blood feeding. **(C)** NOS expression analysis after *Fer+Trf* KD showing that after KD there is no significant change found in brain after blood feeding taking SF brain as a control. Two independent biological replicates were statistically analyzed with the help of one-way ANOVA (Dunnett's multiple comparison test) where asterisks represented *p < 0.05; **p < 0.005; and ***p < 0.0005. (N = number of biological replicates, n= number of mosquitoes dissected for sample collection).

**Table 4: Summary of *in-silico* homology search analysis of newly identified transcripts encoding novel proteins in the mosquito *An. culicifacies***

| S. No. | Protein Name | Length (AA) | Species | Identity (in %) | Role |
|---|---|---|---|---|---|
| **1.** | Hepcidin | 59 | *Mugil incilis* | 100 | Regulation of iron level |
| **2.** | Transferrin 1 | 638 | *Mugil cephalus* | 90.61 | Uptake of iron and iron homeostasis |
| **3.** | Ceruloplasmin | 88 | *Mugil cephalus* | 94.32 | Carrier for copper |

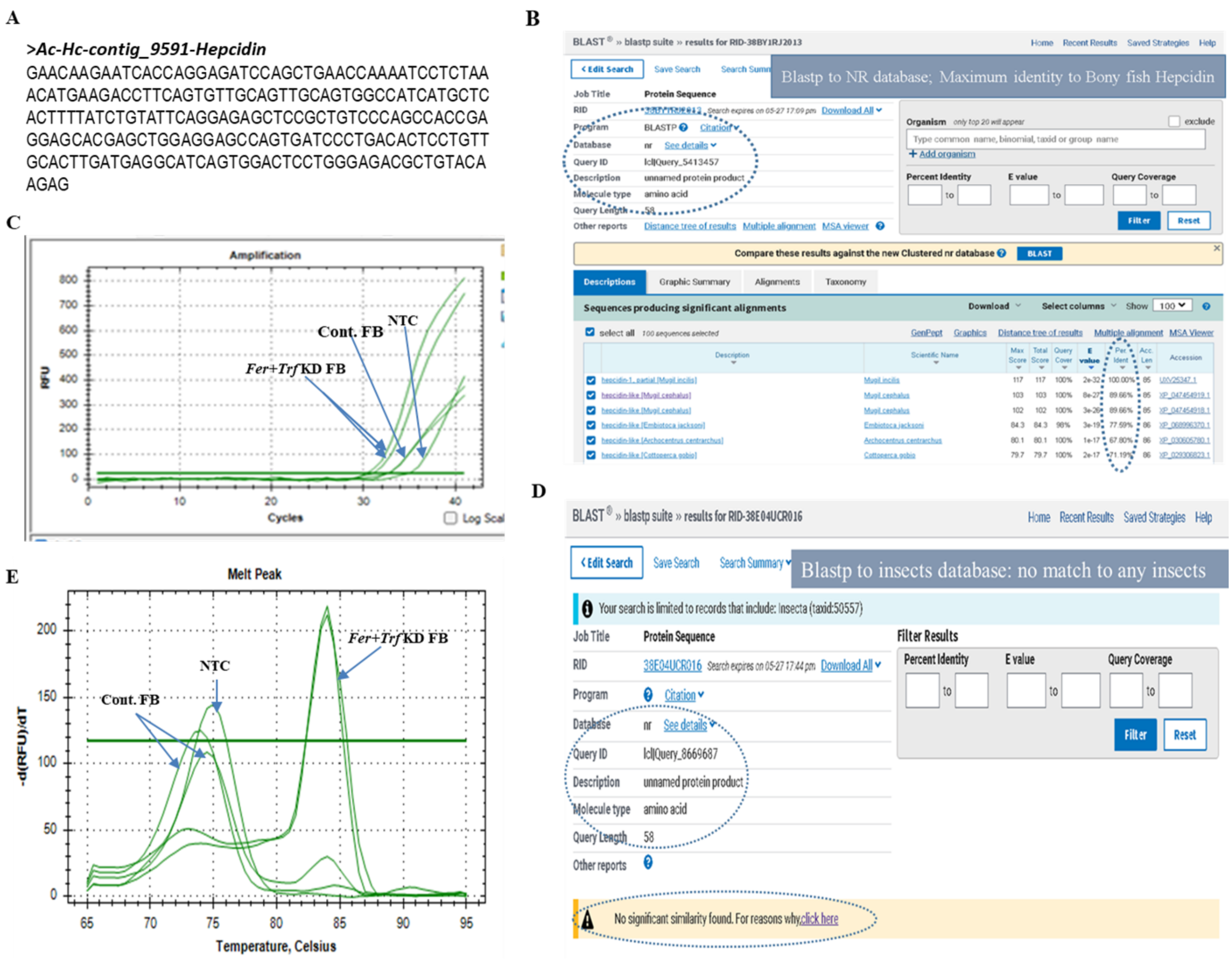


**Fig. 8: Analysis of *Hepcidin*: (A)** Nucleotide sequence of *Hepcidin* **(B)** Online NCBI based BLASTp matrix of mosquito *Hepcidin*, highest identity (100%) to bony fishes **(C)** Amplification curve analysis of qPCR showing that *Fer+Trf* KD start amplification at lower CT value as compared to Control and No-template control. **(D)** Online NCBI-based BLASTp matrix of mosquito *Hepcidin*, showing no similarity to any insect species to the NR database. **(E)** Melting curve analysis of qPCR showing different melting curves of *Fer+Trf* KD, Control, and No-Template Control.

**Table 5:** Iron homeostasis disruption elevates nitrite level in mosquito's brain: Relative change in the nitrite levels pattern measured by absorbance of sugar fed, blood fed and *Fer+Trf* KD mosquito's brain

| Sample | Nitrite detection (µm/ml) |
|---|---|
| SF_Control | 1.03 |
| SF_Silenced | 1.535 |
| BF_Control | 2.187 |
| BF_Silenced | 2.5078 |
